# Protection against *N. gonorrhoeae* induced by OMV-based Meningococcal Vaccines are associated with cross-species directed humoral and cellular immune responses

**DOI:** 10.1101/2024.11.29.626107

**Authors:** Weiyan Zhu, Andreea Waltmann, Marguerite B. Little, Kristie L. Connolly, Kathryn A. Matthias, Keena S. Thomas, Mary C. Gray, Aleksandra E. Sikora, Alison K. Criss, Margaret C. Bash, Andrew N. Macintyre, Ann E. Jerse, Joseph A. Duncan

## Abstract

Limited protective immunologic responses to natural *N. gonorrhoeae* infection and a lack of knowledge about mechanisms of protection have hampered development of an effective vaccine. Recent studies in humans and mice have found meningococcal outer membrane vesicle-containing vaccines (OMV) induce cross species immune responses against gonococci and are associated with protection. The exact mechanisms or how humoral and cellular immunity are related to protection, remain unclear. To study this, we immunized mice with two meningococcal OMV-containing vaccines known to accelerate clearance of *N. gonorrhoeae*, 4CMenB and OMV from an engineered *N. meningitidis* strain lacking major surface antigens PorA, PorB, and Rmp (MC58 ΔABR). We assessed serologic and cellular immune signatures associated with these immunizations and assessed bacterial clearance in the mice using a vaginal/cervical gonococcal infection model. Mice immunized with 4CMenB or MC58 ΔABR demonstrated shortened courses of recovery of vaginal *N. gonorrhoeae* compared to control mice immunized with alum alone. Vaccination with 4CMenB or MC58ΔABR OMV elicited serum and vaginal cross-reactive anti-Ng-OMV antibody responses that were augmented after vaginal challenge with *N. gonorrhoeae*. Further, splenocytes in 4CMenB and MC58 ΔABR immunized mice exhibited elevated cytokine production after restimulation with heterologous *N. gonorrhoeae* OMV when compared to splenocytes from Alum immunized mice. We further tested for correlations between bacterial burden and the measured anti-gonococcal immune responses within each vaccination group and found different immunologic parameters associated with reduced bacterial burden for each vaccine. Our findings suggest the cross-protection against gonococcal infection induced by different meningococcal OMV vaccines is likely multifactorial and mediated by different humoral and cellular immune responses induced by these two vaccines.

## 1. Introduction

*Neisseria gonorrhoeae* is a highly prevalent bacterial sexually transmitted pathogen that can cause both symptomatic or asymptomatic infection of the genital tract and other mucosal surfaces. *N. gonorrhoeae* infection is typically treated with antibiotic therapy, but the pathogen has remarkable capacity for the development of antibiotic resistance. *N. gonorrhoeae* isolates with resistance to all classes of antibiotic used to treat these infections have been identified(1). Vaccines would be the most efficient way to mitigate antimicrobial resistance. Despite the urgent need, the development of *N. gonorrhoeae* vaccines has been hindered by a lack of understanding of how host immunity eliminates this pathogen and provides protection. Recurrent infections with *N. gonorrhoeae* are common, and it is believed that most humans infected with *N. gonorrhoeae* fail to develop protective immunity. Natural immune responses to infection in female mouse models of gonococcal infection are also inadequate to drive protection(2). However, when vaginal immune responses are driven toward Th1 polarization through blockade of endogenous IL-17 polarizing cytokines or through administration of Th1 polarizing cytokines, accelerated clearance of the bacteria has been observed(3–5). The protection observed in these murine studies required intact serologic and cellular immune functions.

For many years, gonorrhea vaccine research efforts struggled after failure of a vaccine against the *N. gonorrhoeae* pilus in field trials suggested phase variable expression and antigenic shift of gonococcal surface antigens could prevent the human host from mounting key protective immune responses(6). In recent years, retrospective case-control studies have found a reduction in the incidence of gonorrhea following vaccination with meningococcal serogroup B OMV containing vaccines. Although the estimated efficacy of these vaccines in preventing gonococcal infection is relatively low, ∼30-40% effectiveness, this epidemiologic evidence of vaccine induced cross-species protective immunity has reinvigorated the gonococcal vaccine field (7–13). Unlike capsular polysaccharide-based meningococcal vaccines, group B meningococcal OMV vaccines target cell surface proteins (14–16). Because *Neisseria gonorrhoeae* and *Neisseria meningitidis* are closely related, sharing between 80 and 90% genome sequence identity, their OMV contain numerous conserved vaccine candidates, including proteins and glycolipids (17–21). In theory, cross-protective immunity could arise through the highly conserved antigens shared by both species. Cross reactive serologic immune responses against *N. gonorrhoeae* proteins in both humans and mice immunized with a licensed OMV-containing vaccine has been reported (22). Additionally, cross species immune reactivity to *N. gonorrhoeae* has been observed in mice immunized with pre-clinical stage OMV based vaccines generated from *N. meningitidis* strains engineered to lack major immunodominant membrane antigens (23). Mice immunized with either a commercial OMV-containing MenB vaccine (4CMenB) through the subcutaneous route or a preclinical MenB OMV vaccine through the intraperitoneal route have been shown to have accelerated clearance of *N. gonorrhoeae* infection but the immunologic mechanism driving this enhanced clearance has not yet been determined. We sought to learn whether the mechanisms of protection against *N. gonorrhoeae* driven by these two different vaccine formulations administered through two different routes could be determined by comparing an analysis of humoral and cellular immune responses to each vaccine along with clearance of *N. gonorrhoeae* after vaginal challenge in vaccinated mice.

## 2. Materials and methods

### 2.1 Bacterial strains and culture conditions

*N. gonorrhoeae* strains F62 and FA1090 A25 were used in mouse intravaginal challenge, assays of serum bactericidal and opsonophagocytic activity assays, and to generate OMV as indicated. Gonococcal strains were cultured on GC base medium (Difco) supplemented with Kellogg’s supplement I under 5% CO2 at 37°C. GC-VNCTS agar [GC agar with vancomycin, colistin, nystatin, trimethoprim (VCNTS supplement; Difco) and 100 μg/ml streptomycin (Sm)] and heart infusion agar (HIA) were used to isolate *N. gonorrhoeae* and facultatively anaerobic commensal flora, respectively, from murine vaginal swabs.

### 2.2 Experimental murine vaccination and infection

All animal experiments were conducted at the Uniformed Services University according to the guidelines of the Association for the Assessment and Accreditation of Laboratory Animal Care using a protocol approved by University’s Institutional Animal Care and Use Committee. Female BALB/c mice [4-5 weeks, Charles River Laboratories (Wilmington, MA)] were housed with autoclaved food, water and bedding and allowed to acclimate to the animal facility for 10 days. Mice were immunized three times with 250 µL of 4CMenB (Bexsero™, Glaxo Smith Kline) administered subcutaneously, or with a 1:1 mixture of 12.5 µg MC58 ΔABR (24) and Alhydrogel (InvivoGen) adjuvant administered intraperitoneally [prepared as previously described (23)]; Alhydrogel alone administered subcutaneously was administered as a control at 3-week intervals. Three weeks after the last immunizations, mice in the diestrus stage of the estrous cycle were identified by vaginal smear and treated with three doses of 0.5 mg of 17β-estradiol every other day, beginning two days prior to inoculation. Mice were also treated with streptomycin and vancomycin to reduce colonization by commensal organisms as described. Mice were then intravaginally challenged in two separate experiments with 1×10^6^ CFU of *N. gonorrhoeae* F62 as previously described (22). Vaginal swabs were quantitatively cultured for *N. gonorrhoeae* on days 1, 3, 5, and 7 post-inoculation as described. Culture results were expressed as CFU/ml of vaginal swab suspension (limit of detection: 20 CFU). The percentage of mice with positive cultures at each time point was plotted using a Kaplan Meier curve and analyzed by the Log Rank test. *N. gonorrhoeae* bacterial recovery and vaginal PMN counting area under the curve (AUC) differences between multiple groups were performed by ordinary one-way ANOVA with Tukey’s multiple comparisons. Correlation analyses between bacterial burden and PMN levels used Pearson Correlation Coefficient. Analyses were performed by GraphPad Prism software (GraphPad Software, Version 9, La Jolla, Calif)

### 2.3 Enzyme-Linked Immunosorbent Assays and Immunoblots

Microtiter plates (384-well) were coated with 80 ng/well of *N. gonorrhoeae* strain F62 OMV, and total IgG, IgG1, IgG2a, IgG3 and IgA antibodies from serum and vaginal lavage were measured by standard ELISA(25) using a BioMek i7 liquid handler and isotype-specific detection antibodies from Southern Biotech. Results were reported as endpoint titer or EC50 of a 4PL curve fit using GraphPad Prism software (GraphPad Software, v9 or higher).

Crude outer membranes from *N. gonorrhoeae* strain F62 were prepared with deoxycholate as described and fractionated by sodium dodecyl sulphate–polyacrylamide gel electrophoresis (SDS-PAGE) and transferred to nitrocellulose membranes. Membranes were blocked with 10% milk in TBS-T and probed with individual mouse serum from vaccinated mice collected post-gonococcal challenge. Blots were developed by incubation with horseradish peroxidase–conjugated anti-mouse IgG secondary antibodies (Bio-Rad), followed by Pierce ECL Western Blotting Substrate (ThermoFisher). Antibody titers or levels, and immunoblot intensity level differences among multiple groups were compared by ordinary one-way ANOVA with Tukey’s multiple comparisons using GraphPad Prism software (GraphPad Software, Version 9, La Jolla, Calif). To assess the change in antibody levels post-gonococcal challenge relative to pre-challenge levels, fold changes were calculated using the formula post ÷ pre and expressed on the log_2_ scale. The mean change in bacterial burden (AUC of log_10_ CFU) with respect to antibody levels and immunoblot intensity was assessed with simple linear regression and 95% confidence intervals using STATA18.

### 2.4 Human Complement Serum Bactericidal Assay

SBA was performed as described (26). In brief, twenty microliters of 5 x 10^4^ CFU/mL bacteria were suspended in 2% bovine serum albumin in Hank’s Balanced Salt Solution containing calcium and magnesium (HBSS; Gibco, Catalog # 14025-092), and incubated with 20µL of heat-inactivated sera pooled from 4–6 immunized mice or control antibodies for 15 minutes at 37°C with 5% CO_2_. The immunoglobulin G/immunoglobulin M–depleted normal human serum (NHS) was used as active complement source and added at 4% final concentration. Samples were mixed and incubated for another 30 minutes. Aliquots were plated on GCB agar and incubated overnight. CFUs of test sera were enumerated after overnight culture at 37°C with 5% CO_2._ Differences in bactericidal titers were assessed by unpaired t and Mann Whitney tests. Analyses were performed by GraphPad Prism software (GraphPad Software, Version 9, La Jolla, Calif)

### 2.5 Opsonophagocytic Killing Assay

OPK assays were performed using fresh human neutrophils as effector phagocytes and C6-depleted pooled normal human serum as a source of complement as previously described (26). Neutrophils were purified from the venous blood of healthy human subjects who provided written consent, following a protocol approved by the University of Virginia Institutional Review Board for Health Sciences Research (#13909). In brief, strain FA1090 bacteria (1000 CFU/well) in HBSS were incubated with 20µL of heat-inactivated sera pooled from 4–6 immunized mice for 30 minutes at 37°C with 5% CO_2._ Sixty µL of C6-depleted pooled NHS (5% final concentration) and 100 µL of 2 x 10^6^/mL neutrophils were added to each well, mixed well and incubated for 2 hours at 37°C with 5% CO_2_. Aliquots of 20 µL of each suspension were plated onto GCB agar. CFU were enumerated and presented as the percentage of bacteria killed. Differences in opsonophagocytic killing were assessed by unpaired t and Mann Whitney tests using GraphPad Prism software (GraphPad Software, Version 9, La Jolla, Calif)

### 2.6 Spleen cell restimulation and multiplex cytokine analysis

To assess the immune response to infection, vaccinated mice were inoculated with *N. gonorrhoeae* F62 (26-29 mice/group). Mice were euthanized at the end date of the experiment using CO_2_ overdose, their spleens were harvested, and a single cell suspension of splenocytes was prepared. Briefly, spleens were crushed and forced through a 70µm filter into T cell Media (RPMI 1640, 10% FBS, 1% HEPES, 1% Pen/Strep, 0.1% 2-ME, 1% 1N NaOH, 1% sodium pyruvate, 1% MEM non-essential amino acids, 2% MEM amino acids). Cells were centrifuged at 400 x g for 10 minutes and the pellets were resuspended in ACK lysing buffer (Thermo Fisher Scientific) for the lysis of red blood cells. Cells were washed twice with T cell media and resuspended in T cells media at the concentration of 1×10^7^ cells/mL. Splenocytes were re-stimulated ex vivo by mixing 100µl of cells with 100 µl of [1 µg/ml] Ng-OMV or media as control in 96-well plate. The plates were incubated at 37°C 5% CO_2_ for 48 hours. Supernatants were harvested and stored at -80°C. Thawed samples were analyzed using the Invitrogen Th1/Th2/Th9/Th17/Th22/regulatory T cell (Treg) cytokine 17-plex mouse ProcartaPlex panel (Thermo Fisher Scientific) to determine levels of cytokines/chemokines. Assays were read on a Luminex 200 or FlexMAP 3D reader, and the background-corrected mean fluorescence intensities were compared to standard curves to calculate pg/mL concentrations of each protein using BioPlex Manager v6.2 (Bio-Rad). Results below the lower limit of quantitation (LLOQ) were imputed as half the LLOQ. Differences between groups were assessed with one-way ANOVA followed by no paring Šidák’s multiple comparison test, with a single polled variance. using GraphPad Prism software (GraphPad Software, Version 9, La Jolla, Calif). The effect of each measured cytokine on the mean change in bacterial burden (AUC of log_10_ CFU) was assessed with simple linear regression with 95% confidence intervals using STAT18.

### 2.7 Principal Coordinate Analysis

Principal Coordinates Analysis (PCoA) was conducted in R to assess sample similarity patterns with respect to antibody and antigen-specific cellular responses. Data were preprocessed using the dplyr package for filtering and transforming, and standardized with clusterSim. The pcoa function in the ape package function computed principal coordinates based on a distance matrix, providing a visual representation of sample similarities in reduced dimensional space. The package ggfortify was used for visualization and graphing purposes. The following variables were considered for the generation of the distance matrix: the pre-gonococcal and post-gonococcal serum immunoglobulin, post-gonococcal challenge and vaginal wash immunoglobulin, and the magnitude of the antigen-specific cellular immune responses (as defined by the log_2_ fold change in secreted cytokines measured by multiplex Luminex when the splenocytes of immunized and challenged mice were co-cultured with gonococcal OMV).

## 3 Results

### 3.1 Vaccination with 4CMenB or MC58 **Δ**ABR shorten bacterial colonization in mice after intravaginal gonococcal challenge

Prior studies have demonstrated that immunization with 4CMenB or MC58 ΔABR OMV leads to a shortened duration of infection and reduced recovery of viable *N. gonorrhoeae* bacteria in a murine genital tract infection model (22, 23). In order to determine whether there were shared mechanisms of enhanced clearance between these two OMV-containing vaccines, BALB/C mice were vaccinated with 3 doses of 4CMenB (administered subcutaneously), MC58 ΔABR OMV (administered intraperitoneally), or Alum (administered subcutaneously) at 3-week intervals and subsequently intravaginally challenged with gonococcal strain F62 three weeks after receiving the last immunization (Fig 1). Bacterial colonization in the genital tract was monitored for 7 days by enumerating viable *N. gonorrhoeae* colonies recovered from vaginal swabbing. 4CMenB- and MC58 ΔABR OMV-vaccinated mice both demonstrated a significant increase in bacterial clearance compared to Alum controls (Fig 2A). At day 7, 34.8% of 4CMenB immunized mice and 64.2% of MC58 ΔABR OMV-immunized mice remained infected with *N. gonorrhoeae* while 88.5% of the Alum-injected control animals continued to have recoverable *N. gonorrhoeae*. As expected, the total bacterial burden, measured as the area under the curve for recovered *N. gonorrhoeae* CFU plotted against time for each individual mouse, was lower in animals with clearance of *N. gonorrhoeae* (Fig. 2B). ΔABR-immunized mice had a slight delay in neutrophil influx with significantly lower vaginal neutrophils on day 5, when compared to Alum and 4CMenB (Fig. 2C). However, there was not a statistically significant difference in the total neutrophil influx during the infection measured as the area under the curve of vaginal neutrophil count plotted against time, (Sup Fig 1A) as measured among the three groups. For mice across all groups, the total neutrophil influx was inversely correlated to the total recovered *N. gonorrhoeae* CFU at the individual mouse level (Sup Fig 2). However, when looking groups of animals based on vaccination status only, 4CMenB-immunized animals showed significant inverse correlation between neutrophil influx and total recovered bacterial burden (Fig. 2D). Combined, both 4CMenB and MC58 ΔABR OMV immunization are associated with increased bacterial clearance in the murine gonorrhea infection model. MC58 ΔABR OMV immunization is associated with delay in vaginal inflammation while clearing bacteria more rapidly than the control group. Interestingly, 4CMenB immunization demonstrated a similar inflammatory response to that seen in control group but was associated with a significant inverse correlation between bacterial burden and neutrophil response, raising the possibility that 4CMenB immunization leads to an immune response that enhances neutrophil based clearance of *N. gonorrhoeae* in these mice.

**Figure 1:**
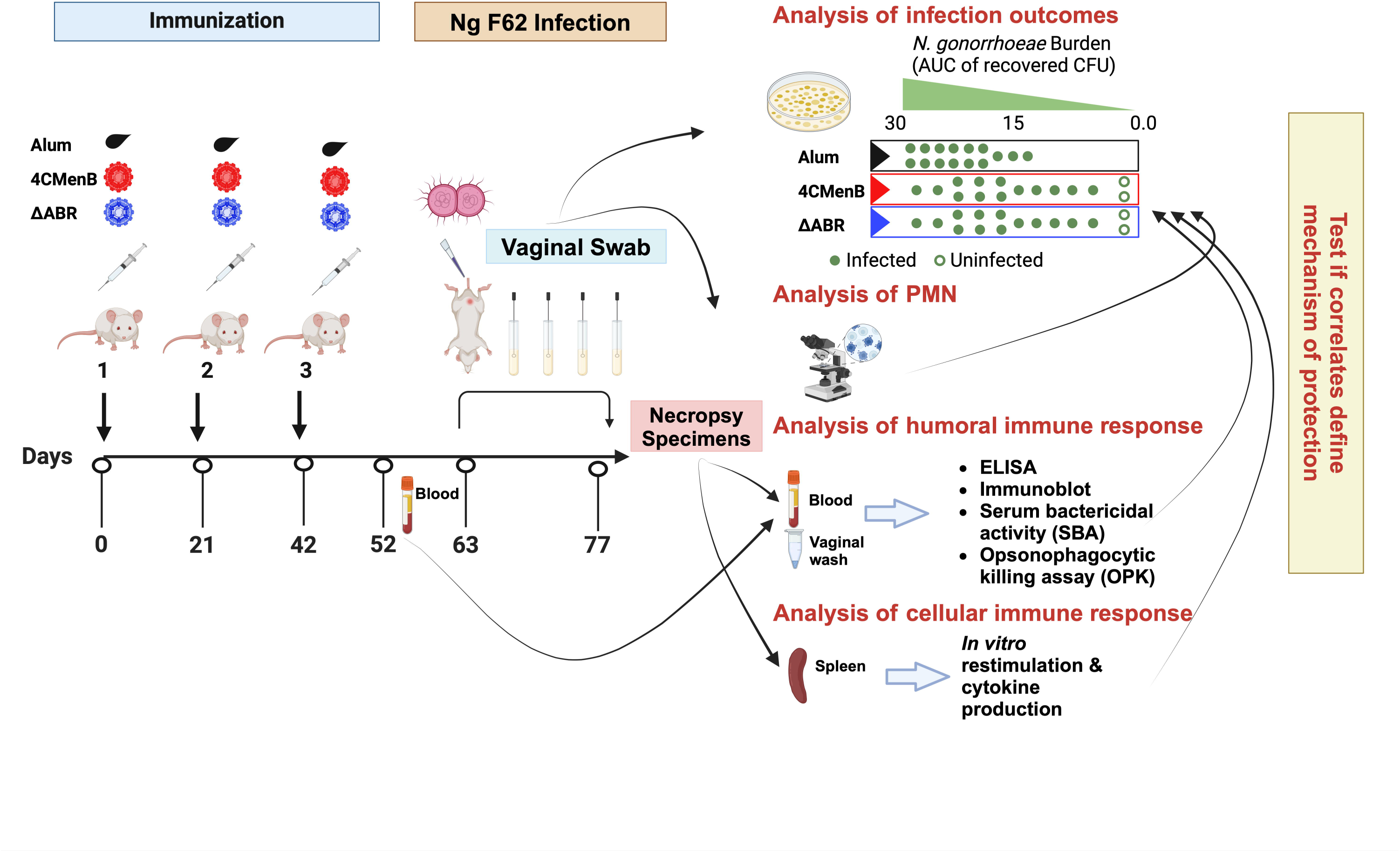
A mouse vaccination followed by vaginal challenge with *N. gonorrhoeae* experimental schema was executed to test whether common immunologic responses to different *N. meningitidis* OMV vaccines that correlated with clearance of infection could be identified. Two identical experiments were performed in which mice were assigned to one of three vaccine groups: Alum (the adjuvant alone control group), MC58 ΔABR OMV vaccine, or 4CMenB vaccine. After the indicated vaccines were administered, an intravaginal challenge with *N gonorrhoeae* strain F62 was conducted and vaginal *N. gonorrhoeae* and PMN influx was assessed over 7 days. Blood, vaginal wash fluid and splenocytes were collected from the mice at the indicated times for the indicated battery of immunologic tests were conducted. The immune responses were tested for correlation to reduced bacterial burden during the period of infection.

**Figure 2:**
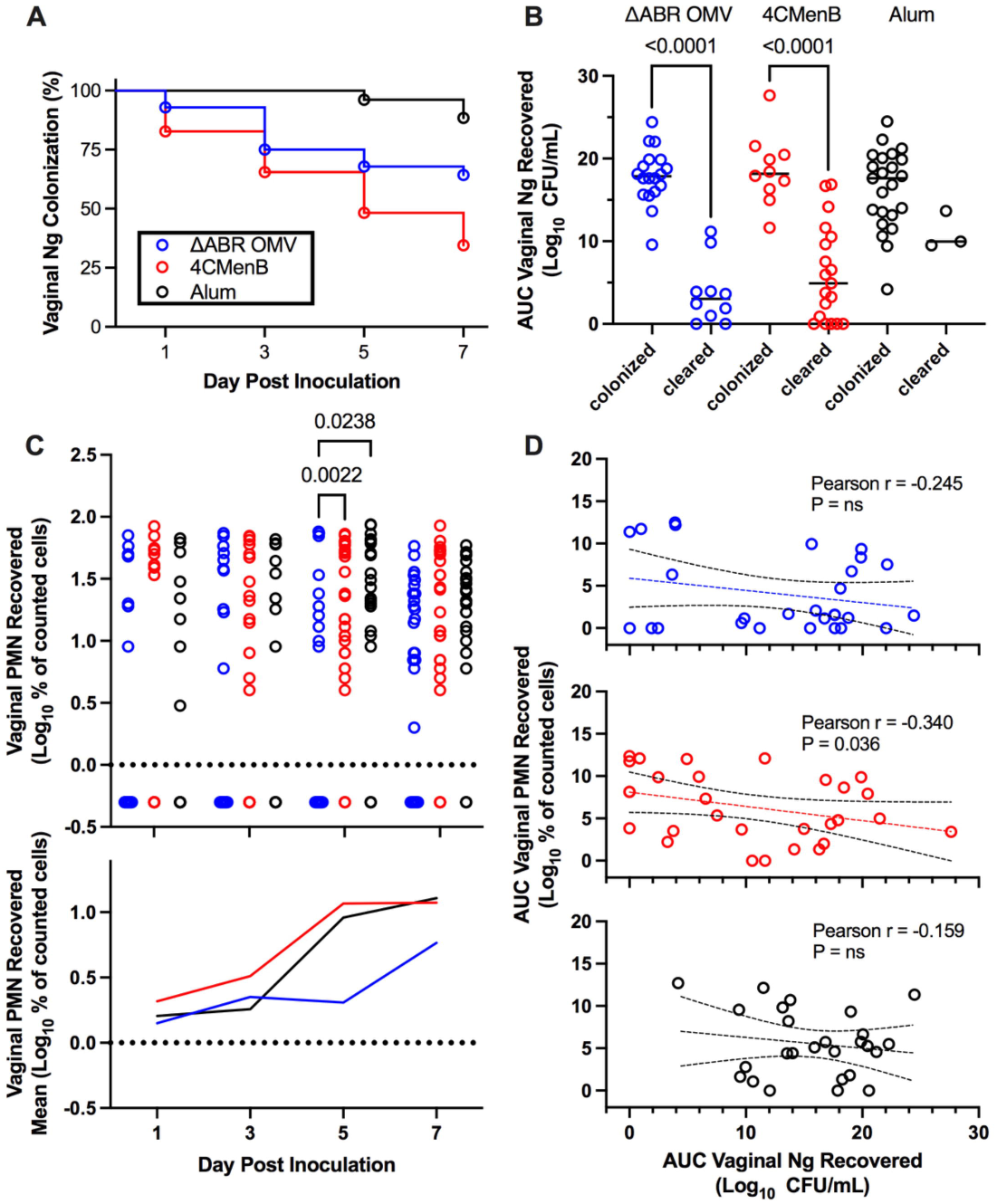
Vaccination with MC58 ΔABR and 4CMenB enhance *N. gonorrhoeae* clearance from lower genital tract in mice. (A) Fraction of mice remaining infected over time with *N. gonorrhoeae* strain F62 after intravaginal inoculation are shown for mice vaccinated with Alum (adjuvant alone control), MC58 ΔABR OMV, and 4CMenB as indicated. Differences between infection persistence were compared using a log-rank (Mentel-Cox) test; *P* were 0.03 or <0.001 when MC58 ΔABR OMV and 4CMenB were compared to the Alum control. (B) Vaginal swab specimens were quantitatively cultured to determine the bacterial burden (number of CFU per milliliter) on days 1,3,5, and 7, the total *N. gonorrhoeae* bacterial burden (AUC vaginal *N. gonorrhoeae* recovered) for the infection course was determined for each individual mouse by taking the area under the curve of the log of the recovered CFU plotted against day post infection. The AUC are plotted for mice in each immunization group and groups are split into those mice that cleared infection by day 7 and those that had persistent infection at day 7. Statistical significance was performed using 2-way ANOVA followed by Bonferroni Post-hoc multiple comparisons to compare immunization groups. (C) Neutrophils (PMN) recovered on vaginal swabs collected on indicated days were quantified and plotted for individual mice in each vaccination group on each day (top panel) and as the mean for the group (bottom panel). (D) The total neutrophil recovery for the course of the infection was determined for each individual mouse by determining the area under the curve of the plotted recovered neutrophils over time and the AUC of Vaginal PMN recovery was plotted against the *N. gonorrhoeae* burden (AUC of Vaginal *N. gonorrhoeae* recovery) for each individual mouse, the Pearson Correlation Coefficient was determined for each immunized group of mice to assess for significant correlations between vaginal PMN and recovered bacteria.

### 3.2 MC58 **Δ**ABR OMV immunization and 4CMenB immunization induce different anti-*N. gonorrhoeae* immunoglobulin responses associated with infection clearance

Antibodies can play important roles in protection against bacterial infection through complement activation, neutralization, adherence blocking, and antibody-dependent cell-mediated cytotoxicity. To compare cross-species directed serological immunity elicited from meningococcal OMV containing vaccines, we collected serum from immunized mice both before and after the *N. gonorrhoeae* challenge and used an ELISA with OMV from *N. gonorrhoeae* strain F62 as a capture antigen to determine the differences in the anti-*N. gonorrhoeae* antibodies between immunized groups. Serum was collected 52 days after the last immunization (pre-challenge serum) and around 10 days after intravaginal challenge with *N. gonorrhoeae* (post-challenge serum). The titers of anti-*N. gonorrhoeae* IgG and IgG1 in sera of mice that received either vaccine were dramatically higher than those found in alum-injected controls, in which almost all animals exhibited titers below the level of detection for the assay (Fig. 3A & B). 4CMenB immunized mice had significantly higher anti-*N. gonorrhoeae* IgG, IgG1 and IgG2a levels than MC58 ΔABR OMV immunized mice. In pre-challenge sera, only 4CMenB-immunized mice demonstrated significantly elevated titers of anti-*N. gonorrhoeae* IgG2a when compared to alum injected animals, though MC58 ΔABR OMV immunized mice did have a trend towards increased *N. gonorrhoeae*-directed IgG2a. (Fig 3A). However, post challenge, both 4CMenB and MC58 ΔABR OMV immunized mice demonstrated increased anti-*N. gonorrhoeae* IgG2a titers when compared to alum immunized animals (Fig 3B). Following vaginal *N. gonorrhoeae* infection, 4CMenB immunized mice demonstrated an increase of anti-*N. gonorrhoeae* specific IgG1 titer and both 4CMenB and MC58 ΔABR OMV immunized mice exhibited increased titers of anti-*N. gonorrhoeae* IgG2a in serum (Fig. 3C).). Overall, as previously shown, immunization with these *N. meningitidis* OMV-containing vaccines elicit anti-*N. gonorrhoeae* serologic responses. However, we now show that the level and IgG subtype responses to immunization differ between vaccines as do the responses to antigen exposure during vaginal infection.

**Figure 3:**
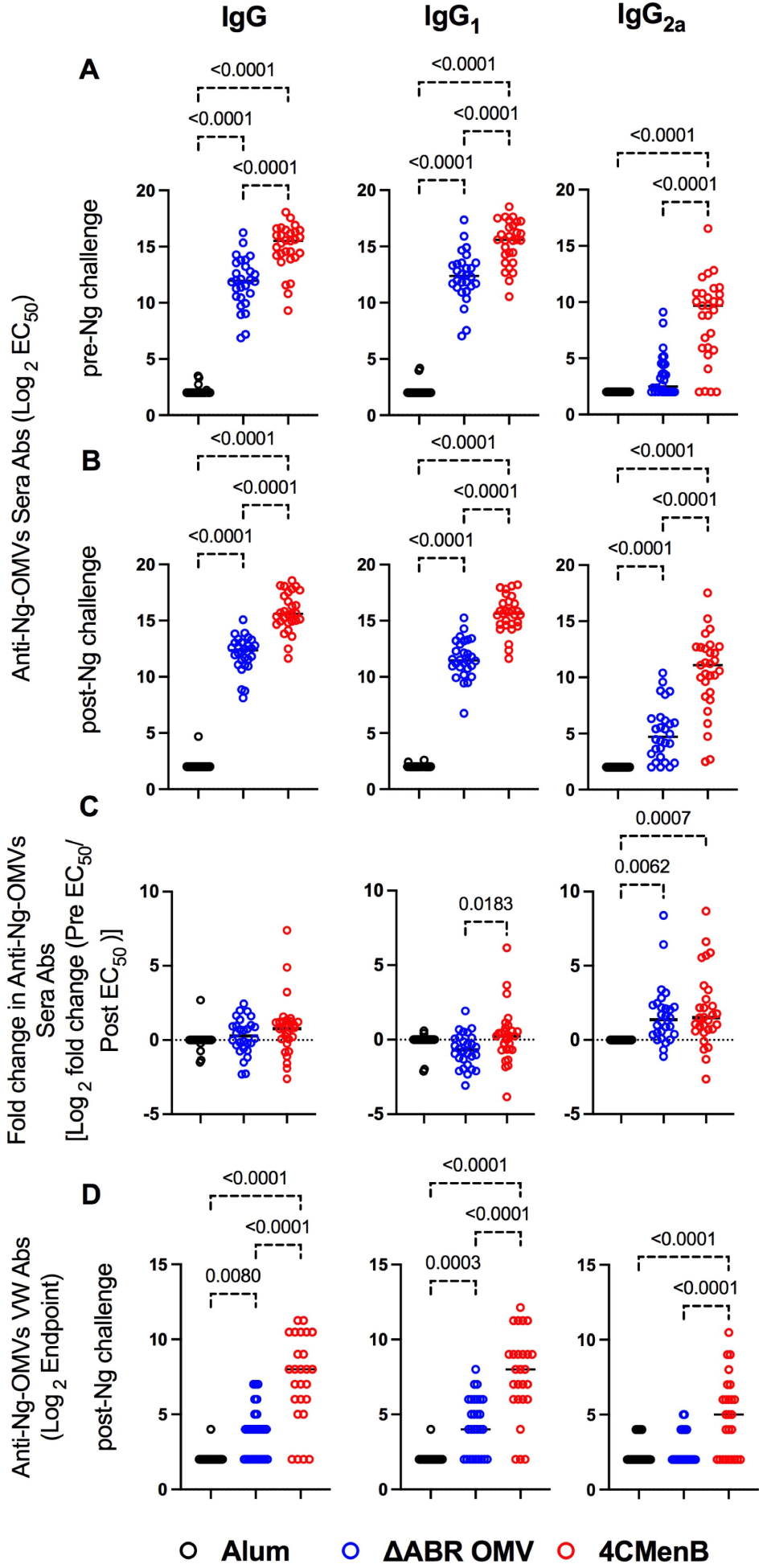
Vaccination with MC58 ΔABR and 4CMenB elicited cross-reactive anti-Ng-OMV antibodies in serum and vaginal fluid. *N. gonorrhoeae* OMV reactive antibody levels were measured by ELISA as described in Materials and Methods in sera collected at day 52, two weeks after final vaccination (A), and at terminal sera collection after *N. gonorrhoeae* challenge (B), and in terminal vaginal wash after *N. gonorrhoeae* challenge (D). The dilution of sera resulting in 50% maximal signal (EC50) and the greatest dilution of vaginal wash fluid resulting in signal above the baseline (Endpoint titer) were determined for total IgG, IgG1 or IgG2a isotype using immunoglobulin subclass specific secondary antibodies. (C) the fold change in each indicated *N. gonorrhoeae* OMV-directed immunoglobulin level between serum collected before challenge and after *N. gonorrhoeae* challenge was also determined. Levels from Alum (black), MC58 ΔABR (blue) and 4CMenB (Red) immunized mice are reported. Data were represented at log scale. Statistical significance was performed using one-way ANOVA followed by no paring Šidák’s multiple comparison test, with a single polled variance.

Because vaginal washing prior to *N. gonorrhoeae* inoculation might impact infection dynamics, we only measured anti-gonococcal vaginal antibody titers at the completion of the experiment. Both vaccine immunization groups demonstrated detectable anti-*N. gonorrhoeae* immunoglobulin levels in their vaginal wash fluid while anti-*N. gonorrhoeae* immunoglobulins were not detected in fluid from alum-immunized mice (Fig 3D). Because all animals had received *N. gonorrhoeae* vaginal inoculations, the lack of anti-gonococcal immunoglobulin in the vaginal washes of alum control mice suggests that while immunization or the combination of immunization and inoculation lead to vaginal anti-*N. gonorrhoeae* antibodies, *N. gonorrhoeae* infection alone was insufficient to induced detectable vaginal anti-gonococcal immunoglobulins.

The relationship between anti-*N. gonorrhoeae* OMV immunoglobulin levels measured by ELISA and the level of *N. gonorrhoeae* recovered from vaginal swabs after intravaginal challenge was examined. In pre-challenge serum, anti-*N. gonorrhoeae* IgG, IgA or specific IgG subtype (IgG1 and IgG2a) titer was not significantly associated with total recovered *N. gonorrhoeae* after challenge in MC58 ΔABR-immunized animals (Fig. 4A). A significant correlation between anti-*N. gonorrhoeae* OMV IgG2a and lower recovered *N. gonorrhoeae* was observed in serum from these animals collected after challenge (Fig 4B). Similarly, the magnitude of change in IgG2a levels after challenge was also correlated with reduced bacterial recovery after MC58 ΔABR immunization (Fig 4C). Despite having higher titers of anti-*N. gonorrhoeae* immunoglobulins than those observed after MC58 ΔABR immunization (Fig. 3), 4CMenB-immunized animals demonstrated no association between these titers and the level of recovered bacteria after challenge (Fig 4A-C). Unexpectedly, anti-*N. gonorrhoeae* immunoglobulins present in vaginal wash fluid after challenge were correlated with increased *N. gonorrhoeae* burden after challenge in MC58 ΔABR immunized animals (Fig 4D).

**Figure 4:**
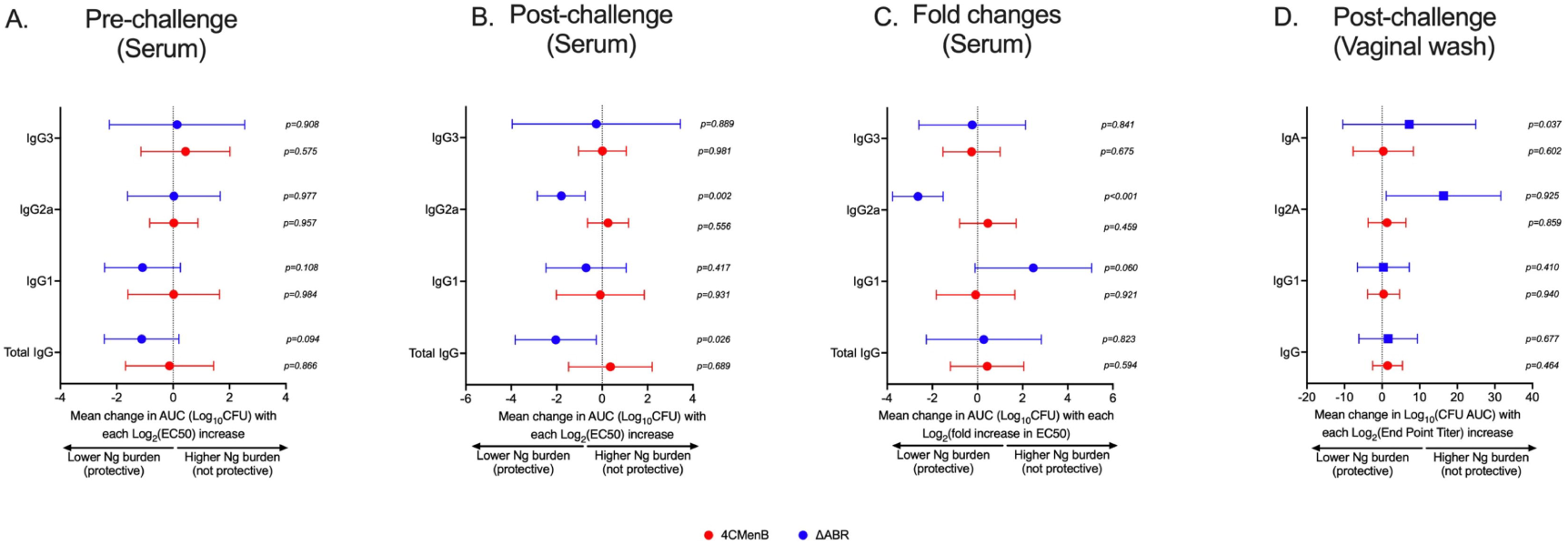
Anti-gonoccoccal IgG2A levels measured by ELISA in serum correlate with reduced gonococcal burden during infection in MC58 ΔABR OMV-immunized mice but not 4CMenB-immunized mice. Mice were immunized with MC58 ΔABR OMV or 4CMenB and subsequently intravaginally challenged with *N. gonorrhoeae* (as described in Figure 1), serum antibody levels of total anti-gonococcal OMV IgG, IgG1, IgG2a and IgG3 were measured by ELISA before and after gonococcal challenge and, levels expressed as log_2_ (EC_50_). The change in levels for each immunoglobulin between levels determined before and after gonococcal challenge were determined as described in Figure 3. Mouse vaginal total anti-gonococcal IgG, IgG1, IgG2a and IgG3 was measured after gonococcal challenge only and measured by ELISA, expressed as log_2_ (endpoint titers). The mean change in bacterial burden (AUC of log_10_ CFU) with respect to serum antibody pre-challenge levels (A), post-challenge levels (B), and fold change(C)) and vaginal wash antibody levels (D) for each mouse was assessed with simple linear regression and 95% confidence intervals are being reported.

We also sought to assess whether vaccination with 4CMenB or MC58 ΔABR elicited antibodies that induced bactericidal antibody activity or promoted human neutrophil opsonophagocytic killing activity against *N. gonorrhoeae*, and whether these functional responses correlated with protection. Sera from mice were pooled to provide adequate volume of specimen to conduct the killing assays. Mice were grouped into pools by the *N. gonorrhoeae* bacterial burden to allow comparisons in these assays between serum from mice that rapidly cleared infection and those in which *N. gonorrhoeae* infection persisted. While both immunizations were associated with increased serum bactericidal activity and opsonophagocytic killing activity when compared to serum from alum immunized animals, the point estimate for SBA titer was higher in MC58 ΔABR immune sera and the point estimate for OPK activity was higher in 4CMenB immune sera (Sup Fig 3A&B). Though the number of pools was insufficient for rigorous statistical testing, the highest titers of these activities were not consistently found in the pools of sera generated from mice with lower bacterial burdens (Sup Fig 3A&B).

SDS-PAGE and immunoblot analysis were used to qualitatively and semi-quantitatively assess the identify of *N. gonorrhoeae* proteins that cross-react with immune sera collected from immunized mice after intravaginal challenge with *N. gonorrhoeae*. Sera from 4CMenB and MC58 ΔABR immunized mice were used individually to probe a membrane carrying proteins from OMV derived from *N. gonorrhoeae* strain F62. Serum from alum immunized animals did not recognize *N. gonorrhoeae* proteins at the dilution used for these experiments (Fig 5A). Serum from each immunized mouse recognized a similar but not completely overlapping set of *N. gonorrhoeae* OMV proteins within each immunization group though the intensity of immunoreactivity against each band had some variation between individual mice (Fig 5A). Antibodies in the sera of MC58 ΔABR immunized mice sera consistently bound to antigens with apparent molecular weights of approximately 85, 66, 55, and 42 kDa, while antibodies in the sera of 4CMenB immunized mice consistently recognized antigens with a molecular weight of approximately 66, 55, 37, and 15 kDa (Fig.5A). The signal intensity for the entire lane and for each of the consistently identified bands were determined using densitometry for each serum specimen used in these immunoblots. The relationship between immunoblot reactivity of each serum specimen and the quantity of *N. gonorrhoeae* recovered during vaginal challenge was explored using linear regression. For ΔABR OMV-immunized mice, the total level of immunoreactivity as well as intensity of the 85 and 66 kDa bands were associated with increased recovered *N. gonorrhoeae*. For 4CMenB-immunized animals, the immunoreactivity against the 37 kDa band was correlated with lower *N. gonorrhoeae* recovery suggesting an association of immunoglobulins directed against this 37 kDa protein with protection from infection. Overall, these serologic studies demonstrate that immunoreactivity against *N. gonorrhoeae* antigens measured by ELISA and immunoblot likely detect antibodies with different properties and ability to provide protection from gonococcal infection.

**Figure 5:**
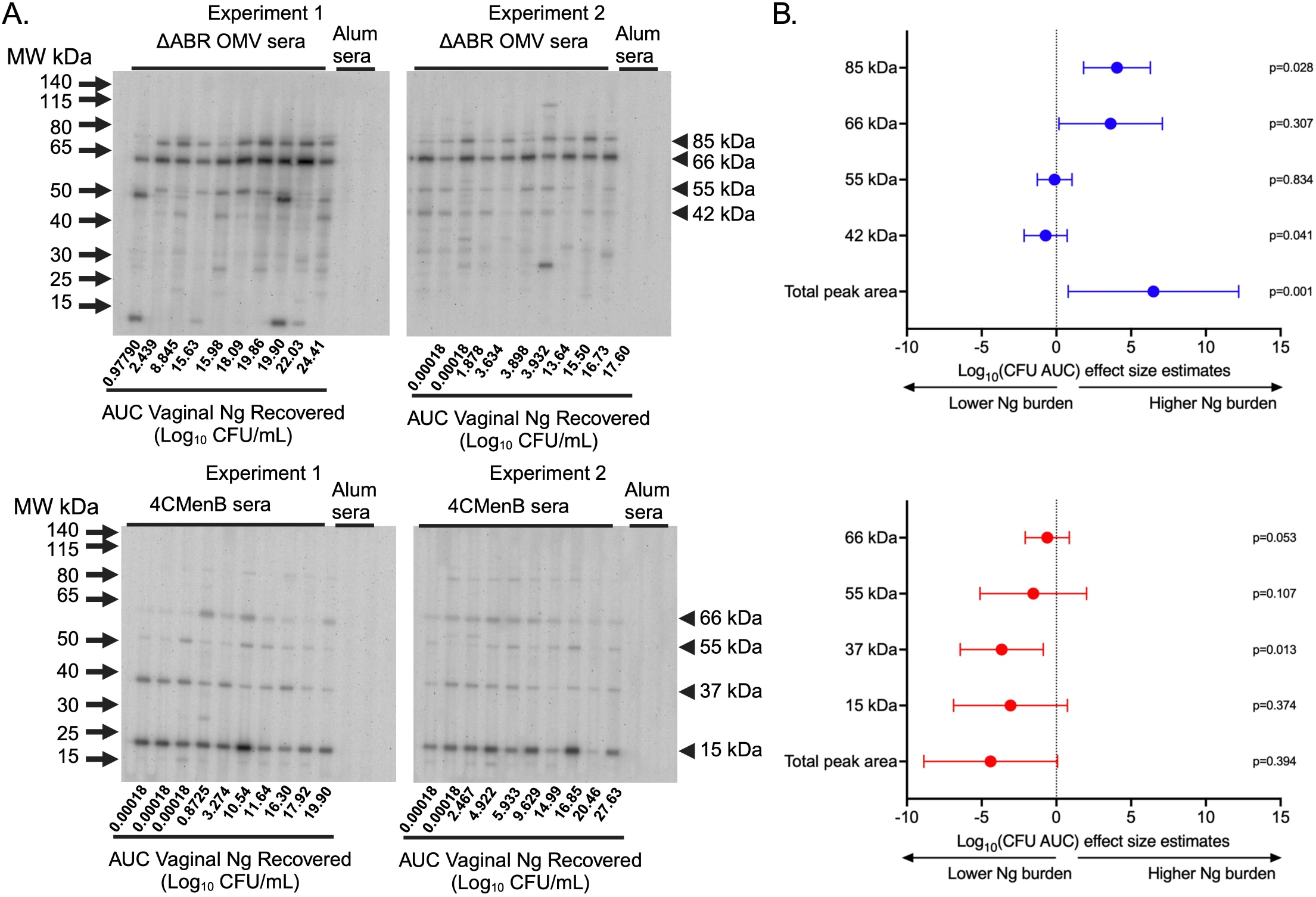
Immunoblot reactivity against *N. gonorrhoeae* OMV proteins correlates with increased *N. gonorrhoeae* burden in MC58 ΔABR OMV-immunized mice and with decreased *N. gonorrhoeae* burden in 4CMenB-immunized mice. (A) Proteins from crude outer membrane vesicle (cOMV) preparations from *N. gonorrhoeae* strain F62 were separated using sodium dodecyl sulfate polyacrylamide gel electrophoresis (SDS-PAGE) and transferred to nitrocellulose. The separated *N. gonorrhoeae* proteins were blotted with individual mouse serum from mice immunized with either MC58 ΔABR OMV (top panel) or 4CMenB (bottom panel) collected after *N. gonorrhoeae* challenge used at 1:1500 dilution in a slot blotting apparatus. Sera from alum-immunized control mice was pooled from each of the two experiments performed and these two pools were used in the indicated slots on each immunoblot as non-vaccine exposed control serum. The order of the analyzed serum specimens from each immunized group was arranged by recovered *N. gonorrhoeae* burden of each mouse and the relative burden of recovered *N. gonorrhoeae* is noted below slot lane for each mouse. The relative antibody reactivity in each slot lane was assessed by densitometric analysis of total peak area for the peak area of each of the 4 most reactive bands common to each immunization type. (B) The relationship between immunoblot signal intensity (assessed by densitometry) and the *N. gonorrhoeae* burden in the vaginal tract of each mouse over the course of infection, expressed as the area under the curve AUC of log_10_ (CFU) recovered from all culture-positive days, was assessed using simple linear regression and 95% confidence intervals.

### 3.3 Vaccination with 4CMenB or MC58 **Δ**ABR generated cell mediated cross-reactive immune responses against Ng

To determine whether 4CMenB and MC58 ΔABR vaccinations activated anti-Ng-OMV specific cellular immune responses, spleens of immunized and *N. gonorrhoeae* challenged mice were harvested, and single-cell suspensions were prepared and cultured in the presence or absence of Ng-OMV as a source of gonococcal antigens. After two days of restimulation, cell supernatants were collected and used to measure production of 12 different cytokines using bead based multianalyte arrays. Baseline splenocyte secretion of all measured cytokines from MC58 ΔABR-immunized and *N. gonorrhoeae* challenged mice was significantly lower than that of Alum control and 4CMen immunized and *N. gonorrhoeae* challenged mice (Sup figure 4). To assess the adaptive immune responses specific to *N. gonorrhoeae* antigens that were specific to immunization, we compared the fold-increase between baseline and stimulation with Ng-OMV (Fig 6A) or 4CMenB (Fig 6 B) for each secreted cytokine in splenocytes harvested from alum, MC58 ΔABR, and 4CMenB immunized mice after *N. gonorrhoeae* challenge was completed. After restimulation with Ng-OMV antigens, splenocytes from 4CMenB and MC58 ΔABR immunized mice exhibited increased IL-2, IL-4 and IL-5 secretion compared to the controls (Fig. 6A). Splenocytes from MC58 ΔABR immunized animals also exhibited enhanced *N.gonorrhoeae* OMV-stimulated IFN-γ, TNF-α, IL-10, and IL-17 when compared to splenocytes from control mice. While splenocytes from MC58 ΔABR immunized animals exhibited stimulation of a broader variety of cytokines than 4CMenB immunized animals in response to *N. gonorrhoeae* OMV, splenocytes from 4CMenB-immunized mice demonstrated robust stimulation of all cytokines measured when stimulated ex vivo with 4CMenB. Linear regression analysis examining relationships between the burden of *N. gonorrhoeae* during infection and splenocyte cytokine production at baseline (Fig 7A), with *N. gonorrhoeae* OMV stimulation (Fig 7B), and with 4CMenB stimulation (Fig 7C) was conducted. Interestingly, the level of 4CMenB stimulated secretion of the immunosuppressive cytokine IL-10 was correlated with increased *N. gonorrhoeae* burden in 4CMenB immunized mice. 4CMenB stimulated IL-1β and IL-2 secretion was also associated with increased *N. gonorrhoeae* burden.

**Figure 6:**
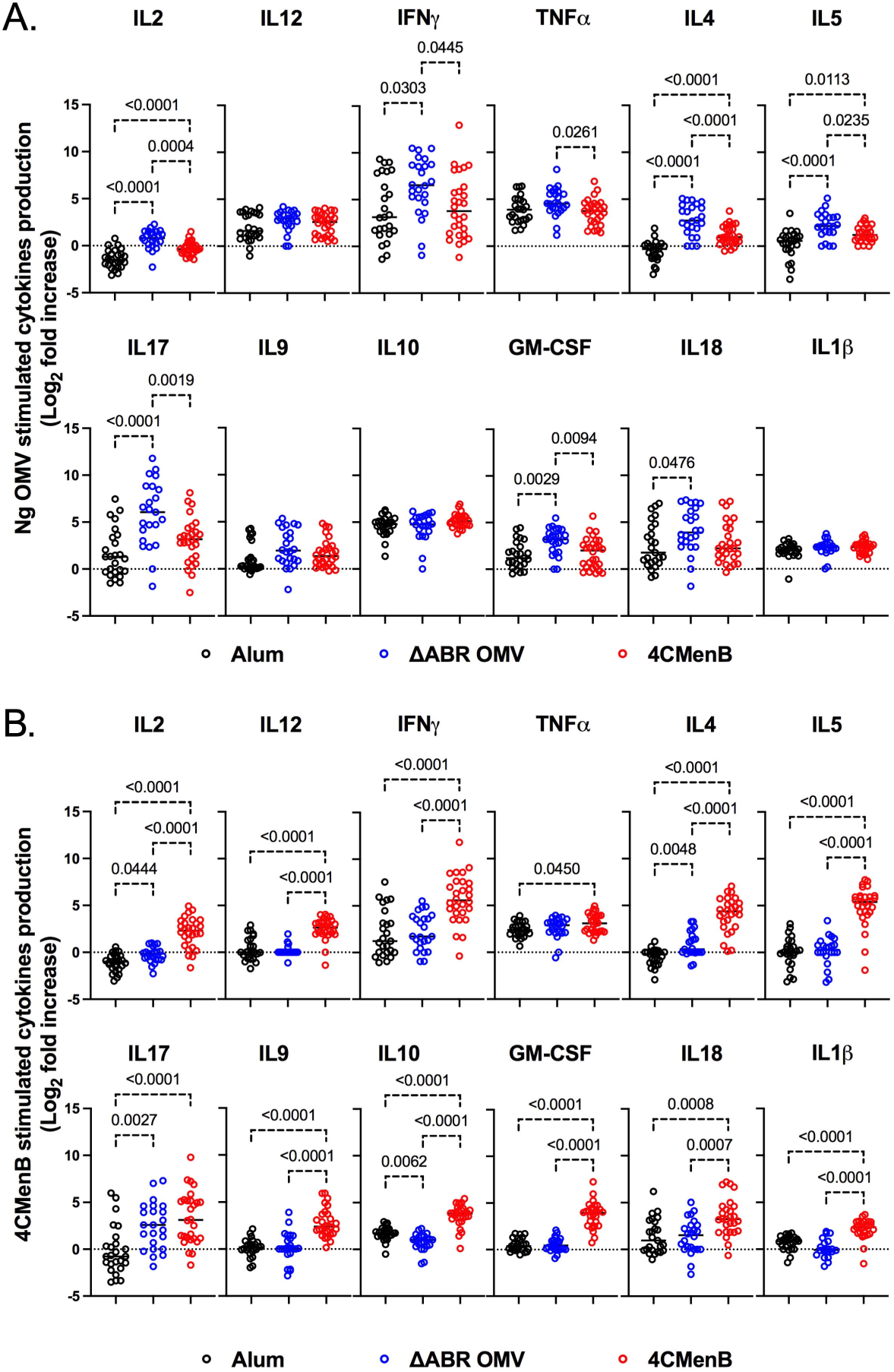
Immunization with MC58 ΔABR OMV and 4CMenB induce N. gonorrhoeae antigen induced cellular immune responses. Splenocytes were collected from mice immunized with Alum, MC58 ΔABR, or 4CmenB and subsequent *N. gonorrhoeae* vaginal challenge. Cells were cultured without stimulation or were stimulated ex vivo with either *N. gonorrhoeae* OMV (A) or 4CMenB (B). After 48 hours, cell culture supernatant was collected and the indicated secreted cytokines were measured using a multiplexed bead array based assay. The antigen-induced response is reported as fold increase in cytokine production in supernatant from antigen stimulated cells and unstimulated cells. Statistical significance was performed using one-way ANOVA followed by no paring Šidák’s multiple comparison test, with a single polled variance.

**Figure 7:**
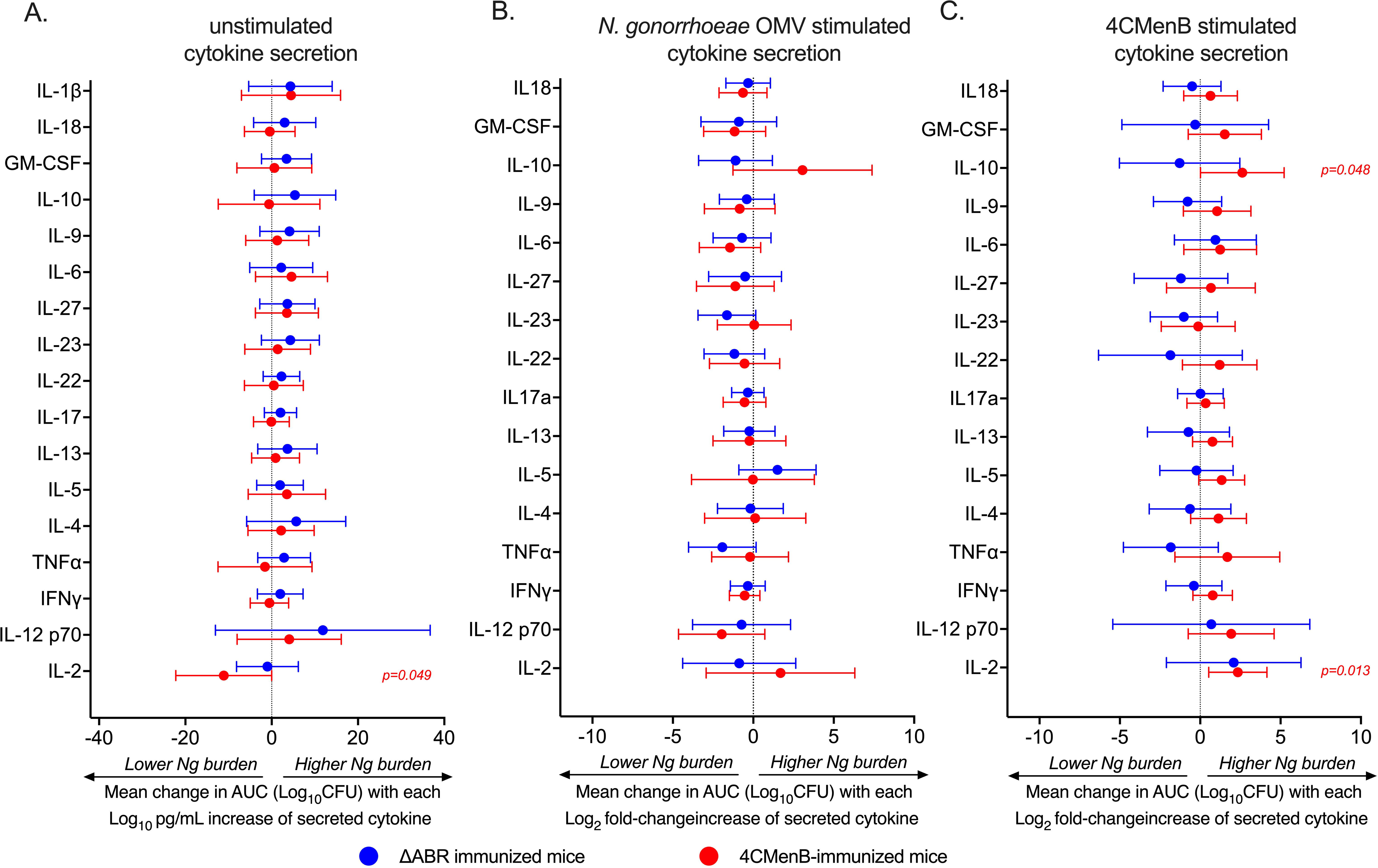
Assessment of the relationship between bacterial burden and cytokine secretion profile of the post-vaccination mouse splenocytes reveals 4CMenB stimulated IL-10 and IL-2 is associated with increased *N. gonorrhoeae* burden in 4CMenB immunized mice. The association between bacterial burden and post-vaccination unstimulated splenocyte cytokine secretion. (A), gonococcal-specific response when splenocytes were stimulated *ex vivo* with *N. gonorrhoeae* OMV (B), and 4CMenB-induced cytokine secretion when splenocytes were stimulated *ex vivo* with 4CMenB as antigen (C) was examined. The effect of each measured cytokine on the mean change in bacterial burden (AUC of log_10_ CFU) was assessed with simple linear regression, the results of which are presented here as forest plots with 95% confidence intervals.

### 3.4 Principal Coordinate Analysis (PCoA) demonstrates differences in profile of immunologic response between vaccine used but no common immune response profile associated with *N. gonorrhoeae* clearance

Our data illustrate that vaccination with 4CMenB and MC58 ΔABR is associated with enhanced gonococcal clearance in a mouse infection model, but immune components (immunological factors) common to both immunizations that were associated with this protection were not identified using simple linear regression analysis to determine the relationship of each parameter with infection intensity. To test whether multiple parameters in combination with one another might yield a common immunologic mechanism for *N. gonorrhoeae* clearance, we conducted unsupervised principal coordinate analysis (Fig 8A). The result showed that mice immunized with Alum, 4CMenB or MC58 ΔABR formed clusters that were distinct from each other, suggesting that different vaccinations exhibited difference in polarization of immune responses, either due to the antigenic make up of each vaccine or due to the route of vaccine administration. When Alum immunized animals were taken out of the PCoA, 4CMenB or MC58 ΔABR continued to cluster based on vaccination group (Fig 8B). However, in both these analyses, there was not clustering of mice that cleared infection separate from mice that did not clear infection. PCoA was also conducted on each group of immunized mice separately to test whether there was within group parameters that were associated with *N. gonorrhoeae* clearance (Fig 8C&D). Neither vaccination group was found to have clustering of mice that cleared or failed to clear *N. gonorrhoeae* infection. Taken together, the PCoA data showed that vaccination yielded distinct immunological profile and that there was not a distinct profile that separated mice based on their clearance of *N. gonorrhoeae* infection.

**Figure 8:**
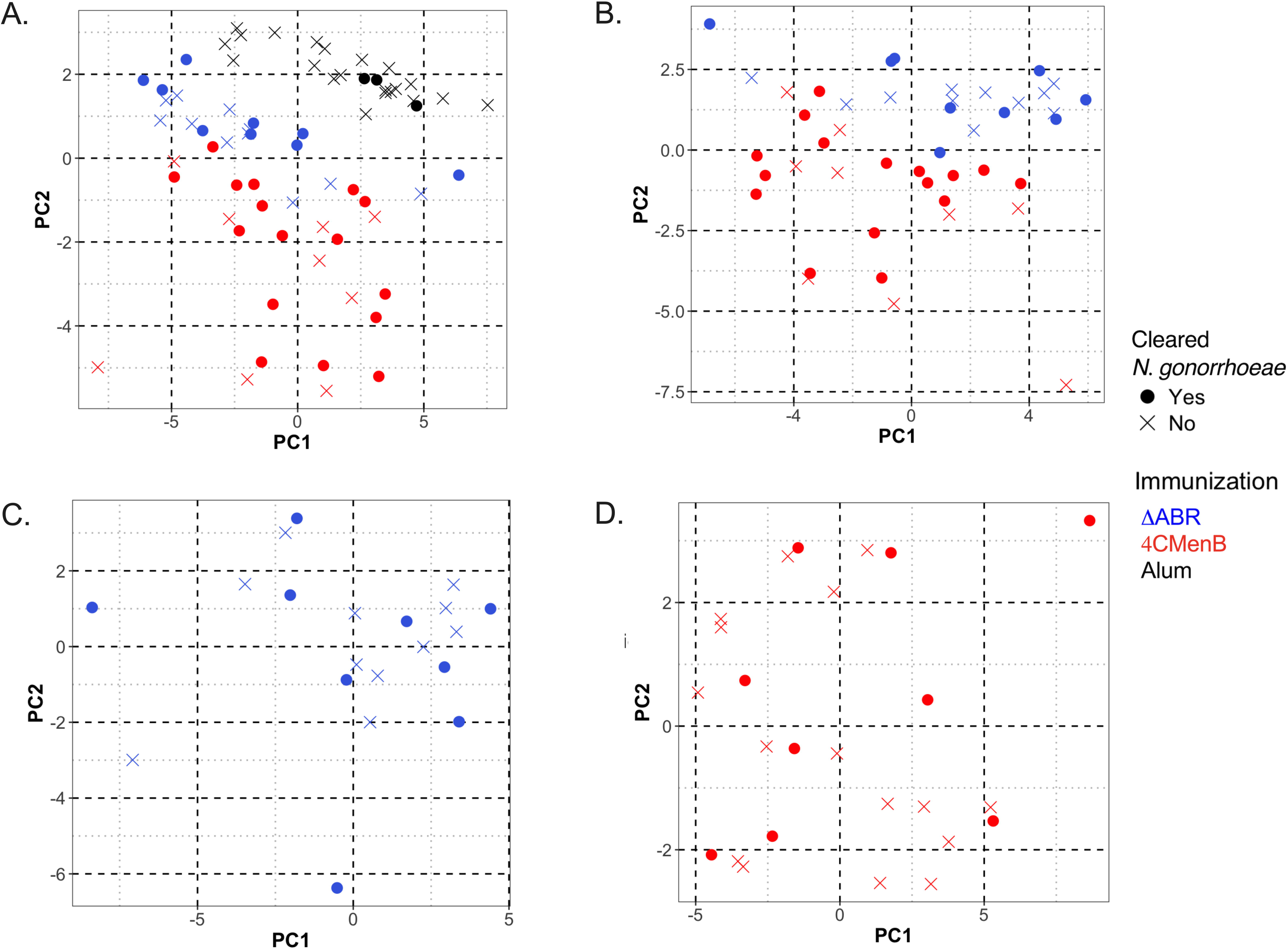
Principal Coordinate Analysis (PCoA) of antibody and cellular immune responses among mice vaccinated with MC58 ΔABR OMV, 4CMenB and Alum control can distinguish immunization type but does not reveal associations between multiple immunologic parameters and clearance of infection. Principal Coordinate Analysis (PCoA) was utilized in R to quantify and visualize the variation in antibody and antigen-specific cellular responses and determine if clustering correlates with vaccination status. Data was preprocessed with dplyr for filtering and transformation, and standardized using clusterSim. The PCoA function in the ape package computed principal coordinates based on a distance matrix, providing a visual representation of sample similarities in reduced dimensional space. The package ggfortify was used to visualize the clustering. The following variables were considered for the generation of the distance matrix: the pre-gonococcal and post-gonococcal serum immunoglobulin, post-gonococcal challenge and vaginal wash immunoglobulin, and the magnitude of the antigen-specific cellular immune responses (as defined by the log2 fold change in secreted cytokines measured by multiplex Luminex when the splenocytes of immunized and *N. gonorrhoeae*-challenged mice were co-cultured with gonococcal outer membrane vesicles, OMV). Data from mice of the three vaccine groups (MC58 ΔABR, 4CMenB, and control mice immunized with Alum adjuvant only) were used. Each point on the PCoA plots corresponds to the immune profile of an individual mouse, with respect to the variables considered. Color coding indicates the three vaccine groups. Open circles denote mice that cleared *N. gonorrhoeae* and closed circles denote mice that were persistently colonized with *N. gonorrhoeae* post-challenge. Clusters indicate relatedness among samples. The differences in immune response were characteristic of the immunization group and could clearly distinguish between the two vaccine groups and Alum control mice (A) and between the two vaccine groups (B), with 66.1% and 62.4% of the total variation in the dataset being explained by the first two principal coordinates, respectively. Within the MC58 ΔABR vaccine group (C) and 4CMenB vaccine group (D), the variation in the multivariable immune response was effectively captured by the first two coordinates (60.8% and 59.1%, respectively) but was not correlated with the ability of the mice to clear infecting gonococci, as evidence by the lack of clustering with respect to gonococcal clearance.

## 4 Discussion

Growing experimental evidence in humans and mice suggests that meningococcal OMV based vaccines provide cross-species protection against the heterologous bacterium, *N. gonorrhoeae*. We have demonstrated that in a mouse model assessing vaccine induced protection to cervicovaginal *N. gonorrhoeae* infection, two different protective vaccines containing *N. meningitidis* OMV induce robust serologic and cellular immune responses against *N. gonorrhoeae* antigens. Lack of detectable antibody to gonococcal antigens is associated with the development of more severe consequences of *N. gonorrhoeae* infection such as salpingitis, suggesting that antibodies may be protective (27–29). Human challenge studies of *N. gonorrhoeae* inoculation in healthy young male volunteers showed a trend toward lower rates of infection in volunteers who developed the strongest antibody responses to previous challenge.(30) To assess the specific antibody responses induced by these *N. meningitidis* OMV containing vaccines that are responsible for protection against *N. gonorrhoeae* infection, we used regression analysis to analyze the correlation between various immunologic indicators and bacterial clearance. We found that *N. gonorrhoeae*-directed cross-specific IgG2 levels in serum measured by ELISA were significantly associated with a reduction in *N. gonorrhoeae* burden in MC58 ΔABR immunized mice while, in those same mice, intensity of serologic responses assessed by immunoblot against *N. gonorrhoeae* OMV was associated with higher *N. gonorrhoeae* burden. Though 4CMenB induced more robust immunoglobulin responses to *N. gonorrhoeae* than MC58 ΔABR OMV, there was not an association of the anti-gonococcal Ig levels measured by ELISA to *N. gonorrhoeae* burden. Two potential explanations for these findings are plausible and not resolved by our experiments. First, there is a maximum protective effect associated with anti-*N. gonorrhoeae* immunoglobulins measured by ELISA in this system. 4CMenB immunization achieved the maximum effect in all immunized animals, while MC58 ΔABR OMV immunization achieved a level of immunoglobulin responses where the maximum protective effect was not reached by most immunized animals. Alternatively, the antigens targeted by IgG2 responses to MC58 ΔABR OMV immunization that are measured by ELISA with *N. gonorrhoeae* OMV have a protective effect while antigens that 4CMenB induces immunoglobulins against are not protective. Interestingly, the intensity of immunoglobulin responses to *N. gonorrhoeae* antigens measured by immunoblot had opposite correlations with bacterial clearance from those observed with immunoglobulins measured by ELISA. In particular, immunoblot intensity of MC58 ΔABR OMV immune sera against gonococcal OMV was associated with increased bacterial burden during *N. gonorrhoeae* cervicovaginal infection and immunoreactivity against a 37 kDa OMV antigen induced by 4CMenB immunization was associated with reduced *N. gonorrhoeae* burden. The identity of this 37 kDa antigen remains to be determined. PorB is the most abundant outer membrane protein in *N. gonorrhoeae* and runs around 37 kDa on SDS-PAGE. When proteins from this region of an SDS PAGE gel are excised and analyzed by mass spectrometry (21, 22), PorB is the most commonly identified protein. However, recombinant PorB based protein vaccines have not been shown to provide protection against *N. gonorrhoeae* in mouse infection models, and whether the 37 kDa protein reactivity in 4CMenB immune sera detected in immunoblot recognizes PorB is an open question(12). Overall, our experiments do not clearly define an immunoglobulin response to immunization common to both OMV-containing vaccines that is associates with protection in mouse infection models.

While serum bactericidal activity (SBA) is accepted as a measurable correlate of protection for vaccines against invasive *N. meningitidis*, the immunologic mechanisms responsible for protection against *N. gonorrhoeae* and a measurable biomarker for such protection remain unknown. Although bactericidal antibodies directed against *N. gonorrhoeae* lipooligosaccharide have been demonstrated to provide protection against *N. gonorrhoeae* in mouse models of cervicovaginal infection and *N. meningitidis* OMV vaccines have been shown to induce bactericidal responses against *N. gonorrhoeae* in mice, our study suggests the bactericidal responses driven by these OMV vaccines are not responsible for the enhanced clearance of *N. gonorrhoeae* observed in vaccinated mice(23, 26, 31). Antibody-mediated, complement-dependent opsonophagocytosis is an accepted correlate of protection for vaccines against *Streptococcus pneumoniae* (32)and has been suggested as a potential protective activity against *Neisseria* infections, this activity also did not have a correlation with *N. gonorrhoeae* clearance in OMV immunized mice(33, 34). Whether there are other antibody-mediated mechanisms of protection that we did not measure remains a possibility. Specifically, whether antigen/immunoglobulin complex mediated activation of cellular FCgamma receptors may induce host responses that aid in bacterial clearance or if there are antibodies that neutralize specific gonococcal virulence factors important in infection was not examined in this study.

The type of cellular immune response required to prevent gonococcal infection in humans is unknown. Using murine *N. gonorrhoeae* infection models, Russell and colleagues reported that blockade of Th17 polarizing cytokines during *N. gonorrhoeae* challenge in mice skewed the resulting immune response towards Th1 polarization and resulted in reduced burden of infection upon rechallenge. Further, when mice were given intravaginal *N. gonorrhoeae* OMV along with microencapsulated IL12, which drives Th1polarization, a protective effect against *N. gonorrhoeae* infection was established in mice(5). Our data show that vaccination with MC58 ΔABR OMV resulted in splenocytes that secreted Th1, Th2, and Th17 related cytokines when stimulated with gonococcal antigens. 4CMenB vaccination led to splenocytes that responded to vaccine restimulation by releasing Th1, Th2, Th17, and Treg related cytokines but primarily Th2 cytokines when cells were restimulated with gonococcal OMV. Surprisingly, given prior reports, the most robust Th1 responses to vaccination were not associated with reduced bacterial burden and neither were the lowest levels of Th17 secretion. In fact, no specific cytokine responses in MC58 ΔABR OMV-immunized animals were strongly associated with reduced bacterial burden. In 4CMenB immunized mice, the highest levels of induced IL-10, a known immunosuppressive cytokine produced by T regulatory cells, correlated with greatest bacterial burden. This finding is consistent with the observation that blockade of IL-10 in mice is also associated with increased clearance of bacteria in the absence of immunization. *N. gonorrhoeae* infection in humans is accompanied by increased localized levels of IL-10. These findings suggest that 4CMenB may induce both protective and non-protective or even detrimental immune responses to *N. gonorrhoeae* antigens, raising the possibility that the partial protection seen in human observational studies could be associated with humans that have poor IL-10 responses to immunization. A combination of immunologic manipulations in mice is needed to assess which responses are mechanistically protective in this model.

While this manuscript was in preparation a manuscript examining cellular immune responses induced by 4CMenB immunization in mice and the relationship of those responses to protection from vaginal challenge with *N. gonorrhoeae* was deposited by Zeppa et al as a preprint manuscript (35). There were minor differences in how assays were conducted and in the analysis, such as Zeppa et al compared results between “Protected” and “Non Protected” groups to test for association with protection while we have utilized recovered bacterial burden as a continuous variable for regression analysis. Despite these differences, our findings reported in this manuscript largely corroborate the lack of association between vaccine induced cytokine secretion and protection from infection in 4CMenB and alum immunized mice. Our studies lack the single cell resolution of immune cell populations generated by flow cytometry based analysis that are reported by Zeppa and colleagues. However, we did observe an association between antigen-stimulated IL-10 production and increased bacterial burden that was not reported by Zeppa and colleagues. Our manuscript also demonstrates cellular immune responses against heterologous gonococcal antigens rather than recall responses against 4CMenB alone, confirming as expected that *N. meningitidis* OMV based vaccines elicit cellular immune responses reactive against gonococcal antigens. This manuscript also extends beyond the reports of Zeppa and colleagues by testing the relationship between vaccine induced immune responses and protection from gonococcal infection for two different *N. meningitidis*-directed vaccines. Overall, our presented work indicates that two OMV based vaccines administered through two different routes, each inducing enhanced clearance of *N. gonorrhoeae,* may achieve that protection via different mechanisms. Our findings support the possibility that there may be multiple mechanistic pathways through which a gonococcal vaccine could achieve protection in humans. Importantly, a determination of the effectiveness of 4CMenB in humans and further correlation of responses to the vaccine to the protected state in humans is paramount to further development of *N. gonorrhoeae* vaccines.

## Conflict of Interest

JAD has a spouse who is employed by GlaxoSmithKline (GSK), the manufacturer of the 4CMenB vaccine, which was utilized in this study. Neither the author’s spouse nor GSK was involved in funding, designing, conducting, analyzing the research reported in this manuscript. JAD acknowledges that there is a potential conflict of interest related to the employment status of his spouse with GSK and attests that the research conducted and reported in this manuscript is free of any bias that might be associated with the commercial goals of GSK. The remainder of the authors declare that they have no commercial or financial relationships that could be construed as a potential conflict of interest.

## Author Contributions

WZ-Writing-original draft, methodology, investigation; AW-Writing-original draft and review and editing, formal analysis; MBL-Writing-original draft, methodology, investigation; KLC-methodology, investigation; KAM-writing-review and editing, methodology, investigation; KST-writing-review and editing, methodology, investigation; MCG - methodology, investigation; AES-resources, writing-review and editing; AKC-resources, writing-review and editing, methodology, supervision; MCB-resources, writing-review and editing, methodology, supervision; ANM-resources, writing-review and editing, methodology, supervision; AEJ–conceptualization, funding acquisition, supervision; JAD-conceptualization, funding acquisition, supervision, writing-review and editing

## Funding

This work was supported by the National Institutes of Health (U19-AI113170 to A. E. J.; R01-AI117235 to A.E.S.). Biomarker profiling was performed in the Regional Biocontainment Laboratory (RBL) at Duke University, which received partial support for construction and renovation from the National Institutes of Health (UC6-AI058607 and G20-AI167200), and facility support from the National Institutes of Health (UC7-AI180254).

## Acknowledgment

None.

## Data Availability Statement

Datasets are available on request: The raw data supporting the conclusions of this article will be made available by the authors, without undue reservation. Further inquiries can be directed to the corresponding author.

## References

1. Ohnishi M, Golparian D, Shimuta K, Saika T, Hoshina S, Iwasaku K, et al. Is Neisseria gonorrhoeae initiating a future era of untreatable gonorrhea?: detailed characterization of the first strain with high-level resistance to ceftriaxone. Antimicrob Agents Chemother. 2011;55(7):3538–45. DOI: 10.1128/aac.00325-11

2. Lovett A, Duncan JA. Human Immune Responses and the Natural History of Neisseria gonorrhoeae Infection. Front Immunol. 2018;9:3187. DOI: 10.3389/fimmu.2018.03187

3. Liu Y, Egilmez NK, Russell MW. Enhancement of adaptive immunity to Neisseria gonorrhoeae by local intravaginal administration of microencapsulated interleukin 12. J Infect Dis. 2013;208(11):1821–9. DOI: 10.1093/infdis/jit354

4. Liu Y, Islam EA, Jarvis GA, Gray-Owen SD, Russell MW. Neisseria gonorrhoeae selectively suppresses the development of Th1 and Th2 cells, and enhances Th17 cell responses, through TGF-beta-dependent mechanisms. Mucosal Immunol. 2012;5(3):320–31. DOI: 10.1038/mi.2012.12

5. Liu Y, Hammer LA, Liu W, Hobbs MM, Zielke RA, Sikora AE, et al. Experimental vaccine induces Th1-driven immune responses and resistance to Neisseria gonorrhoeae infection in a murine model. Mucosal Immunol. 2017;10(6):1594–608. DOI: 10.1038/mi.2017.11

6. Jerse AE, Bash MC, Russell MW. Vaccines against gonorrhea: current status and future challenges. Vaccine. 2014;32(14):1579–87. DOI: 10.1016/j.vaccine.2013.08.067

7. Abbasi J. New Hope for a Gonorrhea Vaccine. Jama. 2017;318(10):894–5. DOI: 10.1001/jama.2017.11037

8. Petousis-Harris H, Paynter J, Morgan J, Saxton P, McArdle B, Goodyear-Smith F, et al. Effectiveness of a group B outer membrane vesicle meningococcal vaccine against gonorrhoea in New Zealand: a retrospective case-control study. Lancet. 2017;390(10102):1603–10. DOI: 10.1016/s0140-6736(17)31449-6

9. Seib KL. Gonorrhoea vaccines: a step in the right direction. Lancet. 2017;390(10102):1567–9. DOI: 10.1016/s0140-6736(17)31605-7

10. Wang B, Giles L, Andraweera P, McMillan M, Almond S, Beazley R, et al. Effectiveness and impact of the 4CMenB vaccine against invasive serogroup B meningococcal disease and gonorrhoea in an infant, child, and adolescent programme: an observational cohort and case-control study. Lancet Infect Dis. 2022;22(7):1011–20. DOI: 10.1016/s1473-3099(21)00754-4

11. Abara WE, Bernstein KT, Lewis FMT, Schillinger JA, Feemster K, Pathela P, et al. Effectiveness of a serogroup B outer membrane vesicle meningococcal vaccine against gonorrhoea: a retrospective observational study. Lancet Infect Dis. 2022;22(7):1021–9. DOI: 10.1016/s1473-3099(21)00812-4

12. Zhu W, Thomas CE, Chen CJ, Van Dam CN, Johnston RE, Davis NL, et al. Comparison of immune responses to gonococcal PorB delivered as outer membrane vesicles, recombinant protein, or Venezuelan equine encephalitis virus replicon particles. Infect Immun. 2005;73(11):7558–68. DOI: 10.1128/IAI.73.11.7558-7568.2005

13. Bruxvoort KJ, Lewnard JA, Chen LH, Tseng HF, Chang J, Veltman J, et al. Prevention of Neisseria gonorrhoeae With Meningococcal B Vaccine: A Matched Cohort Study in Southern California. Clin Infect Dis. 2023;76(3):e1341–e9. DOI: 10.1093/cid/ciac436

14. Danzig L. Meningococcal vaccines. Pediatr Infect Dis J. 2004;23(12 Suppl):S285-92. DOI:

15. Girard MP, Preziosi MP, Aguado MT, Kieny MP. A review of vaccine research and development: meningococcal disease. Vaccine. 2006;24(22):4692–700. DOI: 10.1016/j.vaccine.2006.03.034

16. Rüggeberg JU, Pollard AJ. Meningococcal vaccines. Paediatr Drugs. 2004;6(4):251–66. DOI: 10.2165/00148581-200406040-00004

17. Perrin A, Bonacorsi S, Carbonnelle E, Talibi D, Dessen P, Nassif X, et al. Comparative genomics identifies the genetic islands that distinguish Neisseria meningitidis, the agent of cerebrospinal meningitis, from other Neisseria species. Infect Immun. 2002;70(12):7063–72. DOI: 10.1128/iai.70.12.7063-7072.2002

18. Hadad R, Jacobsson S, Pizza M, Rappuoli R, Fredlund H, Olcén P, et al. Novel meningococcal 4CMenB vaccine antigens - prevalence and polymorphisms of the encoding genes in Neisseria gonorrhoeae. Apmis. 2012;120(9):750–60. DOI: 10.1111/j.1600-0463.2012.02903.x

19. Tinsley CR, Nassif X. Analysis of the genetic differences between Neisseria meningitidis and Neisseria gonorrhoeae: two closely related bacteria expressing two different pathogenicities. Proc Natl Acad Sci U S A. 1996;93(20):11109–14. DOI: 10.1073/pnas.93.20.11109

20. Findlow J, Borrow R, Snape MD, Dawson T, Holland A, John TM, et al. Multicenter, open-label, randomized phase II controlled trial of an investigational recombinant Meningococcal serogroup B vaccine with and without outer membrane vesicles, administered in infancy. Clin Infect Dis. 2010;51(10):1127–37. DOI: 10.1086/656741

21. Zielke RA, Wierzbicki IH, Weber JV, Gafken PR, Sikora AE. Quantitative proteomics of the Neisseria gonorrhoeae cell envelope and membrane vesicles for the discovery of potential therapeutic targets. Mol Cell Proteomics. 2014;13(5):1299–317. DOI: 10.1074/mcp.M113.029538

22. Leduc I, Connolly KL, Begum A, Underwood K, Darnell S, Shafer WM, et al. The serogroup B meningococcal outer membrane vesicle-based vaccine 4CMenB induces cross-species protection against Neisseria gonorrhoeae. PLoS Pathog. 2020;16(12):e1008602. DOI: 10.1371/journal.ppat.1008602

23. Matthias KA, Connolly KL, Begum AA, Jerse AE, Macintyre AN, Sempowski GD, et al. Meningococcal Detoxified Outer Membrane Vesicle Vaccines Enhance Gonococcal Clearance in a Murine Infection Model. J Infect Dis. 2022;225(4):650–60. DOI: 10.1093/infdis/jiab450

24. Matthias KA, Reveille A, Connolly KL, Jerse AE, Gao YS, Bash MC. Deletion of major porins from meningococcal outer membrane vesicle vaccines enhances reactivity against heterologous serogroup B Neisseria meningitidis strains. Vaccine. 2020;38(10):2396–405. DOI: 10.1016/j.vaccine.2020.01.038

25. Samo M, Choudhary NR, Riebe KJ, Shterev I, Staats HF, Sempowski GD, et al. Immunization with the Haemophilus ducreyi trimeric autotransporter adhesin DsrA with alum, CpG or imiquimod generates a persistent humoral immune response that recognizes the bacterial surface. Vaccine. 2016;34(9):1193–200. DOI: 10.1016/j.vaccine.2016.01.024

26. Gray MC, Thomas KS, Lamb ER, Werner LM, Connolly KL, Jerse AE, et al. Evaluating vaccine-elicited antibody activities against Neisseria gonorrhoeae: cross-protective responses elicited by the 4CMenB meningococcal vaccine. Infect Immun. 2023;91(12):e0030923. DOI: 10.1128/iai.00309-23

27. Hicks CB, Boslego JW, Brandt B. Evidence of serum antibodies to Neisseria gonorrhoeae before gonococcal infection. J Infect Dis. 1987;155(6):1276–81. DOI: 10.1093/infdis/155.6.1276

28. Plummer FA, Simonsen JN, Chubb H, Slaney L, Kimata J, Bosire M, et al. Epidemiologic evidence for the development of serovar-specific immunity after gonococcal infection. J Clin Invest. 1989;83(5):1472–6. DOI: 10.1172/JCI114040

29. Plummer FA, Chubb H, Simonsen JN, Bosire M, Slaney L, Nagelkerke NJ, et al. Antibodies to opacity proteins (Opa) correlate with a reduced risk of gonococcal salpingitis. J Clin Invest. 1994;93(4):1748–55. DOI: 10.1172/JCI117159

30. Schmidt KA, Schneider H, Lindstrom JA, Boslego JW, Warren RA, Van de Verg L, et al. Experimental gonococcal urethritis and reinfection with homologous gonococci in male volunteers. Sex Transm Dis. 2001;28(10):555–64. DOI: 10.1097/00007435-200110000-00001

31. Gulati S, Shaughnessy J, Ram S, Rice PA. Targeting Lipooligosaccharide (LOS) for a Gonococcal Vaccine. Front Immunol. 2019;10:321. DOI: 10.3389/fimmu.2019.00321

32. Kuroda E, Koizumi Y, Piao Z, Nakayama H, Tomono K, Oishi K, et al. Establishment of a modified opsonophagocytic killing assay for anti-pneumococcal surface protein A antibody. J Microbiol Methods. 2023;212:106804. DOI: 10.1016/j.mimet.2023.106804

33. Granoff DM. Relative importance of complement-mediated bactericidal and opsonic activity for protection against meningococcal disease. Vaccine. 2009;27 Suppl 2(Suppl 2):B117-25. DOI: 10.1016/j.vaccine.2009.04.066

34. Humphries HE, Brookes C, Allen L, Kuisma E, Gorringe A, Taylor S. Seroprevalence of Antibody-Mediated, Complement-Dependent Opsonophagocytic Activity against Neisseria meningitidis Serogroup B in England. Clin Vaccine Immunol. 2015;22(5):503–9. DOI: 10.1128/cvi.00100-15

35. Zeppa JJ, Fegan JE, Maiello P, Islam EA, Lee IS, Pham C, et al. Meningococcal vaccine Bexsero elicits a robust cellular immune response that targets but is not consistently protective against Neisseria gonorrhoeae during murine vaginal infection. bioRxiv. 2024. DOI: 10.1101/2024.09.08.611931

