## supplemental figures for "Protection against *N. gonorrhoeae* induced by OMV-based Meningococcal Vaccines are associated with cross-species directed humoral and cellular immune responses"

**Supplemental Figures and Figure Legends**

**Supplemental Figure 1:**

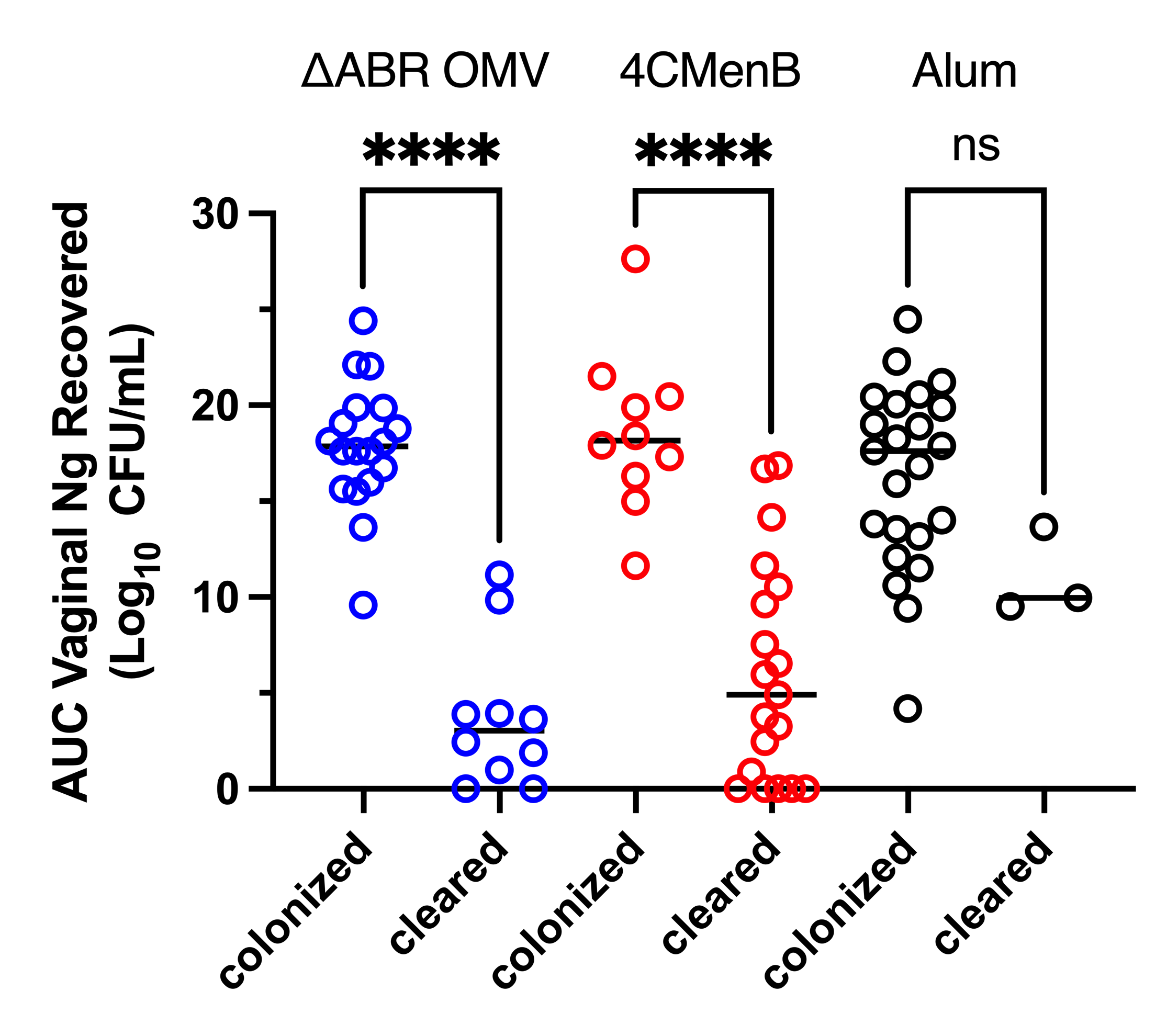

**Supplemental Figure 1: Total vaginal neutrophil influx during *N. gonorrhoeae* challenge in mice was not impacted by vaccination group.** Neutrophils (PMN) recovered on vaginal swabs collected on days 1,3, 5, and 7 were quantified and plotted for individual mice in each vaccination group. The total neutrophil influx for the course of the infection was determined for each individual mouse by determining the area under the curve of the plotted recovered neutrophils over time and plotted for mice from each vaccination group (Alum-black, MC58 ΔABR - blue, and 4CmenB -red), mean neutrophil influx for each group was compared with one-way ANOVA followed by Tukey’s multiple comparisons and no significant difference between groups was identified.

**Supplemental Figure 2:**

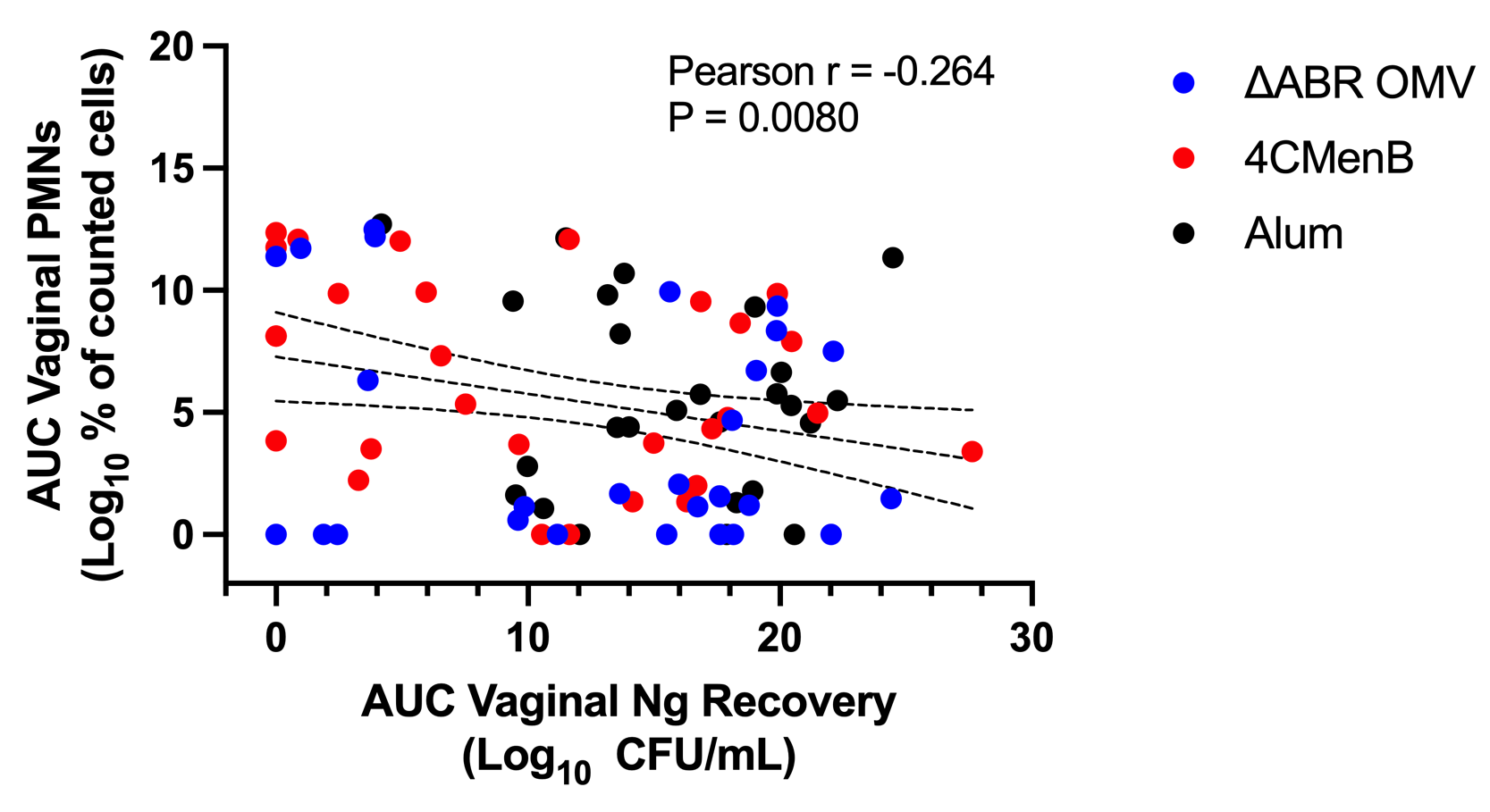

**Supplemental Figure 2:** T**he total neutrophil influx was inversely correlated to the total recovered *N. gonorrhoeae* CFU at the individual mouse level.** The total neutrophil recovery for the course of the infection, expressed as the area under the curve AUC of log_10_ (% of counted cells) recovered vaginal PMN over time, was plotted against the *N. gonorrhoeae* burden (AUC of log_10_ (CFU)) for each individual mouse. The Pearson Correlation Coefficient was determined for all mice to assess for significant correlations between vaginal PMN and recovered bacteria.

**Supplemental Figure 3:**

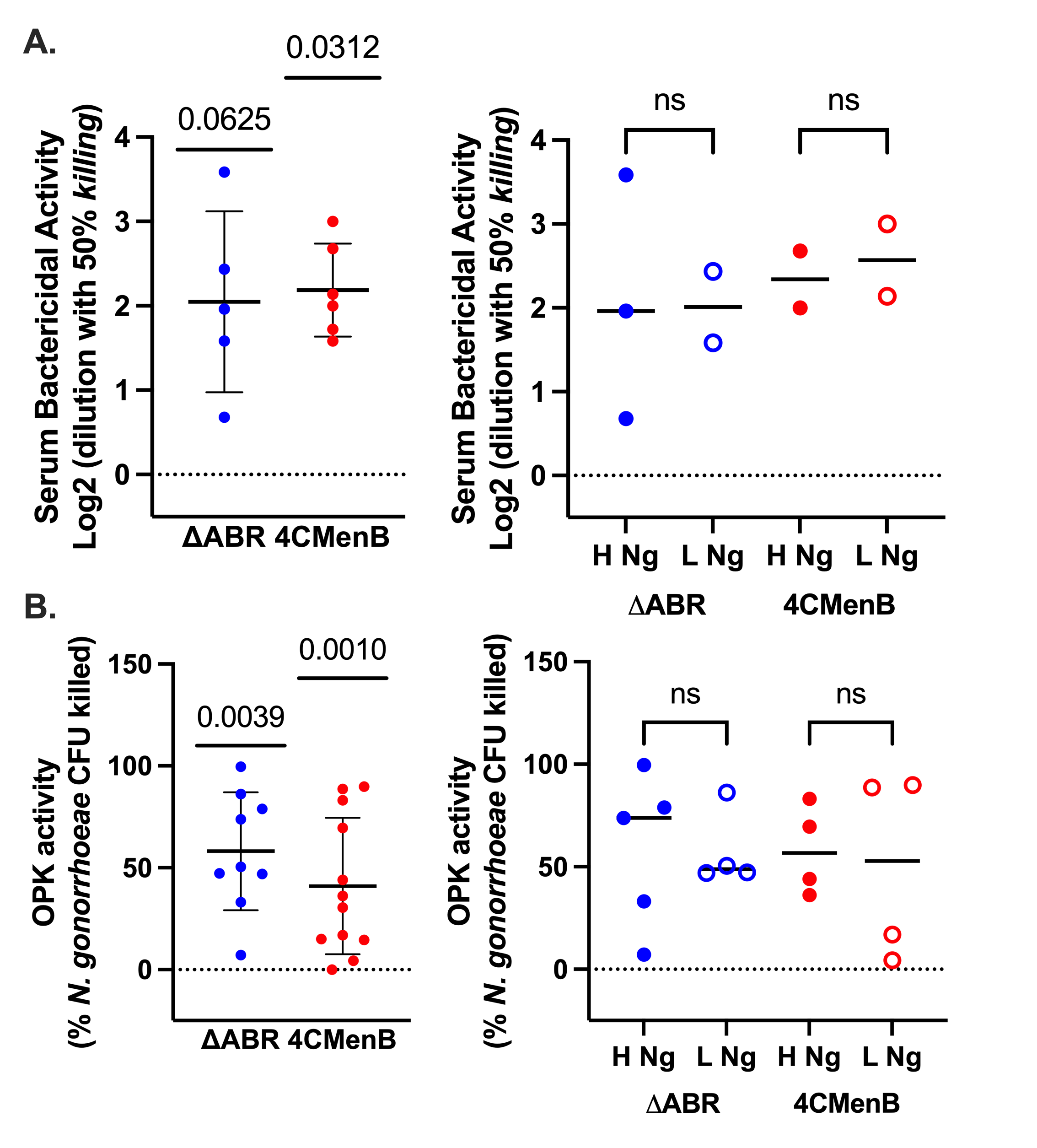

**Supplement Figure 3**: **Both** **4CMenB (red) and MC58 ΔABR (blue) vaccination were associated with increased bactericidal or opsonophagocytic activities compared to Alum controls.** Serum bactericidal activity (**A**) and Opsonophagocytic killing assays (**B**) against *N. gonorrhoeae* was measured from pooled mice sera grouped by level of *N. gonorrhoeae* bacterial recovered during infection. The left panel shows activity of each pool reported as the normalized to pooled serum from alum immunized animals with significance difference from the control sera tested with one sample Wilcoxan test. The right panel shows normalized activity from pooled sera from mice with the highest *N. gonorrhoeae* recovery and those with the lowest recovery during their challenge for each immunization group. Results from the high *N. gonorrhoeae* recovery pools and low *N. gonorrhoeae* recovery pools are compared using one way ANOVA with Sidak correction for multiple comparisons.

**Supplemental Figure 4:**

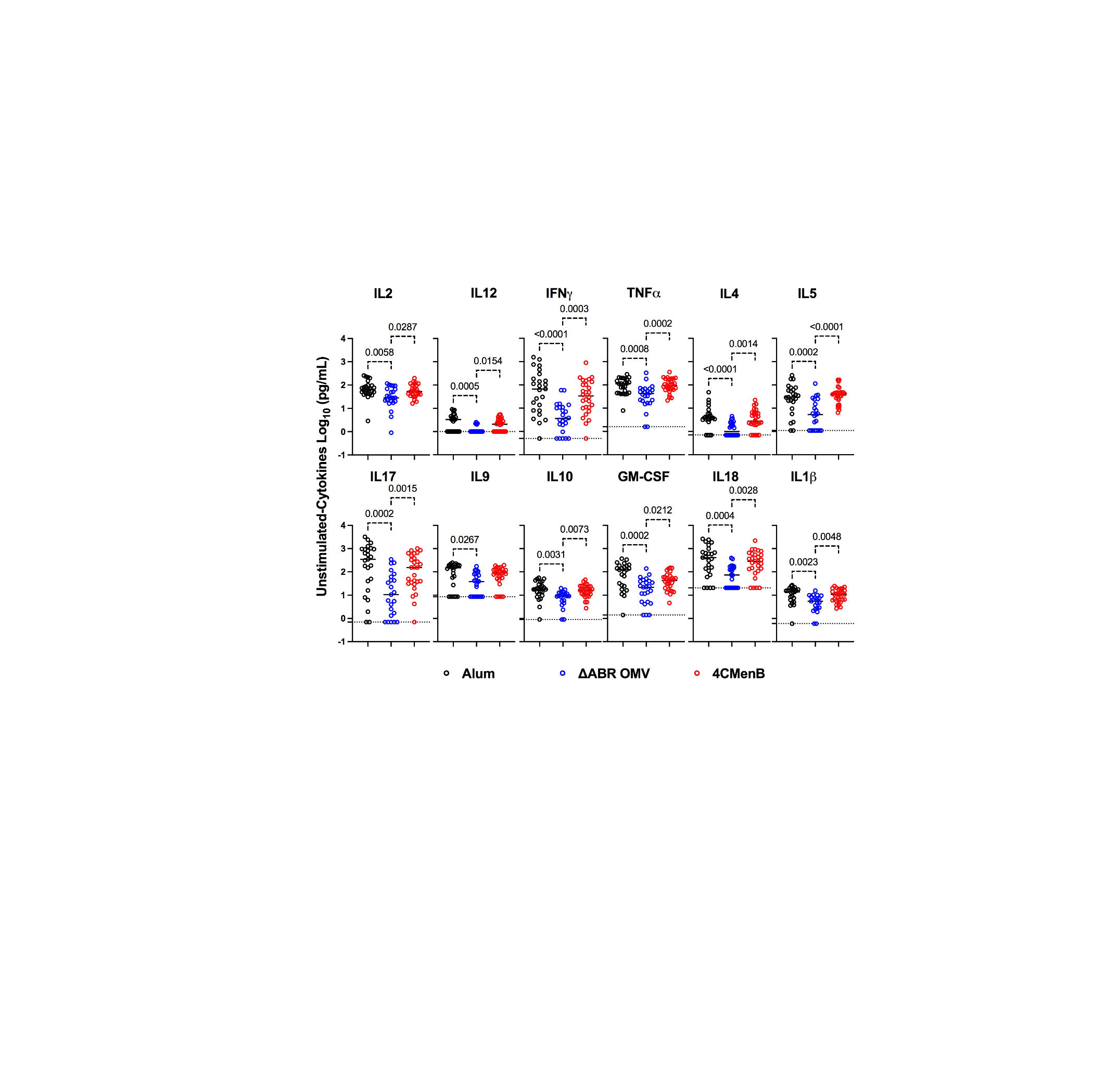

**Supplement Figure 4**: Comparison of unstimulated (baseline) secreted cytokine/chemokine in culture supernatant from cultured splenocytes from Alum (black), MC58 ΔABR (blue) and 4CMenB (red) immunized mice. Data are shown with symbols showing measured level of the indicated cytoine from splenocytes fmor and individual mouse; horizontal bar represents the group mean value. Statistical significance was determined using ordinary one-way ANOVA with Tukey’s multiple comparisons, showing p-values only for significant changes. Created in BioRender. Zhu, W. (2024) <https://BioRender.com/i91g020>
